# Identifying Brain Network Topology Changes in Task Processes and Psychiatric Disorders

**DOI:** 10.1101/866855

**Authors:** Paria Rezaeinia, Kim Fairley, Piya Pal, François G. Meyer, R. McKell Carter

**Affiliations:** Department of Electrical and Computer Engineering, University of California San Diego, San Diego, U.S.A.; Department of Economics, Leiden University, Leiden, The Netherlands; Institute of Cognitive Science, University of Colorado Boulder, Boulder, U.S.A.; Department of Applied Mathematics, University of Colorado Boulder, Boulder, U.S.A.; Department of Electrical, Computer and Energy Engineering, University of Colorado Boulder, Boulder, U.S.A.; Department of Psychology and Neuroscience, University of Colorado Boulder, Boulder, U.S.A.

**Keywords:** fMRI, functional connectivity, random walk, hitting time

## Abstract

**ABSTRACT:** A central goal in neuroscience is to understand how dynamic networks of neural activity produce effective representations of the world. Advances in the theory of graph measures raise the possibility of elucidating network topologies central to the construction of these representations. We leverage a result from the description of lollipop graphs to identify an iconic network topology in functional magnetic resonance imaging data and characterize changes to those networks during task performance and in populations diagnosed with psychiatric disorders. During task performance, we find that task-relevant subnetworks change topology, becoming more integrated by increasing connectivity throughout cortex. Analysis of resting-state connectivity in clinical populations shows a similar pattern of subnetwork topology changes; resting-scans becoming less default-like with more integrated sensory paths. The study of brain network topologies and their relationship to cognitive models of information processing raises new opportunities for understanding brain function and its disorders.

**AUTHOR SUMMARY:** Our mental lives are made up of a series of predictions about the world calculated by our brains. The calculations that produce these predictions are a result of how areas in our brain interact. Measures based on graph representations can make it clear what information can be combined and therefore help us better understand the computations the brain is performing. We make use of cutting-edge techniques that overcome a number of previous limitations to identify specific shapes in the functional brain network. These shapes are similar to hierarchical processing streams which play a fundamental role in cognitive neuroscience. The importance of these structures and the technique is highlighted by how they change under different task constraints and in individuals diagnosed with psychiatric disorders.

## INTRODUCTION

How do we link dynamic changes in functional brain structure to the processing of information? Brain activity organizes into stable networks that vary in strength and change with task demands Greicius, Krasnow, Reiss, and Menon (2003); Smith et al. (2009).

Because of its ease of implementation and relatively low cost, the analysis of resting functional magnetic resonance imaging (rfMRI) data Raichle et al. (2001) in particular has had a tremendous impact, leading to several large-scale public initiatives like the Human Connectome Project (HCP) Essen et al. (2013). One of the most promising methods used to study rfMRI activation has been to construct network models of functional connectivity between areas of the brain E. T. Bullmore and Bassett (2011); Goñi et al. (2014); van den Heuvel and Pol (2010). These models are characterized by network measures like efficiency Fornito, Zalesky, and Bullmore (2016) and have been applied to a wide variety of challenges including the study of psychiatric disorders (for review, see Avena-Koenigsberger, Misic, and Sporns (2017)). Improving our ability to interpret the meaning of these measures for brain processing would have tremendous impact.

To improve our ability to interpret network models of brain connectivity, we seek measures of topology that can be related to models of cognitive information processing. The study of the relationship between brain network topology and function has been accelerating and is key to explaining dynamic information processing in health and disease Stiso and Bassett (2018). To better understand how information is processed in a dynamic context, it is necessary to link specific brain-network topologies to cognitively meaningful information processing structures. Network analysis of brain data typically involves descriptions of an inferred network. Here, we instead describe brain connections as stochastic processes (in our case, using a random walk), avoiding the constraints of a specific network model and instead describing general properties of brain functional connectivity in a given mental state. This improved description of brain connectivity can then be used to link results from graph theory to network topologies common in cognitive models of the brain. As a first step, we utilize a result from the theory of graph measures, which establishes that isolated chains of nodes produce maximally long random walks between points on the graph. In particular, a lollipop graph consists of a set of fully connected nodes attached to a chain of linearly-connected nodes. In a random walk on a lollipop graph, the number of hops required to reach the tail of the lollipop stick is greater than for other topological structures Brightwell and Winkler (1990). We target extremely long random walks between brain areas as a measure of the presence, and relative isolation, of linear chains of nodes. We note that this topology is similar to that found in hierarchical processing streams, a structure important in cognitive models. We hypothesize that those brain areas which take a long time to reach in a random walk are often situated in such an information processing topology.

We focus on the tails of random-walk network connectivity distributions to address the following four key questions. (i) How does the relative isolation of a linear chain of nodes change the distribution of connectivity in a synthetic network? (ii) Are there subnetworks in resting-state cortex that have properties similar to a linear chain of nodes? (iii) How are linear-chain subnetworks changed by task demands? (iv) Does the characterization of network topology have value in understanding and diagnosing psychiatric disorders?

## MATERIALS AND METHODS

### Hitting-time functional connectivity model

One common approach to find the connectivity matrix of a brain network is to threshold the Pearson correlation matrix to obtain the adjacency matrix for the network. Although this method is very simple, it has some shortcomings that might cause inaccuracy in the results. One challenge is that the Pearson correlation coefficient does not account for latent variables, which might result in a high correlation among two regions that are not directly connected. In addition, the choice of threshold is arbitrary, creating interpretation and generalization issues. To overcome these challenges, we integrated the following changes into a standard network analysis pipeline for neuroimaging. First, to compensate for latent variables, we use the partial correlation Smith et al. (2011) to find the connectivity matrix. Let *ρ_ij_* represent the partial correlation between *x_i_* and *x_j_* (the BOLD time series associated with regions *i* and *j*, respectively). Therefore, we use a weighted brain functional network with adjacency matrix *A* = [*ρ_ij_*]. The degree of node *i* is 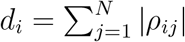. Second, we normalize edge strength using self loops that preserve the overall connectivity of each node relative to others. Third, we characterize the network using the hitting time, a random-walk measure that reflects the expected number of edges that need to be crossed to transition from one node to another. We next describe the edge strength and hitting time approaches in detail.

### Edge Strength Normalization

For a random walk, the probability transition matrix is *P* = [*p_ij_*], where *p_ij_* is defined as:

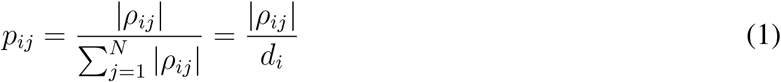

The major drawback of this definition is that it fails to distinguish a strongly connected from a weakly connected node. Consider a network with 5 nodes (a, b, c, d, e) and 6 edges. Suppose that all edges connected to node a have weight 0.9. And, suppose that node b is connected to the same nodes as node a, but with edges with weight 0.1 (Fig. 1A). Applying equation (1), both nodes a and b will have the same transition probabilities, and therefore, the same relative connectivity.

**Figure 1:**
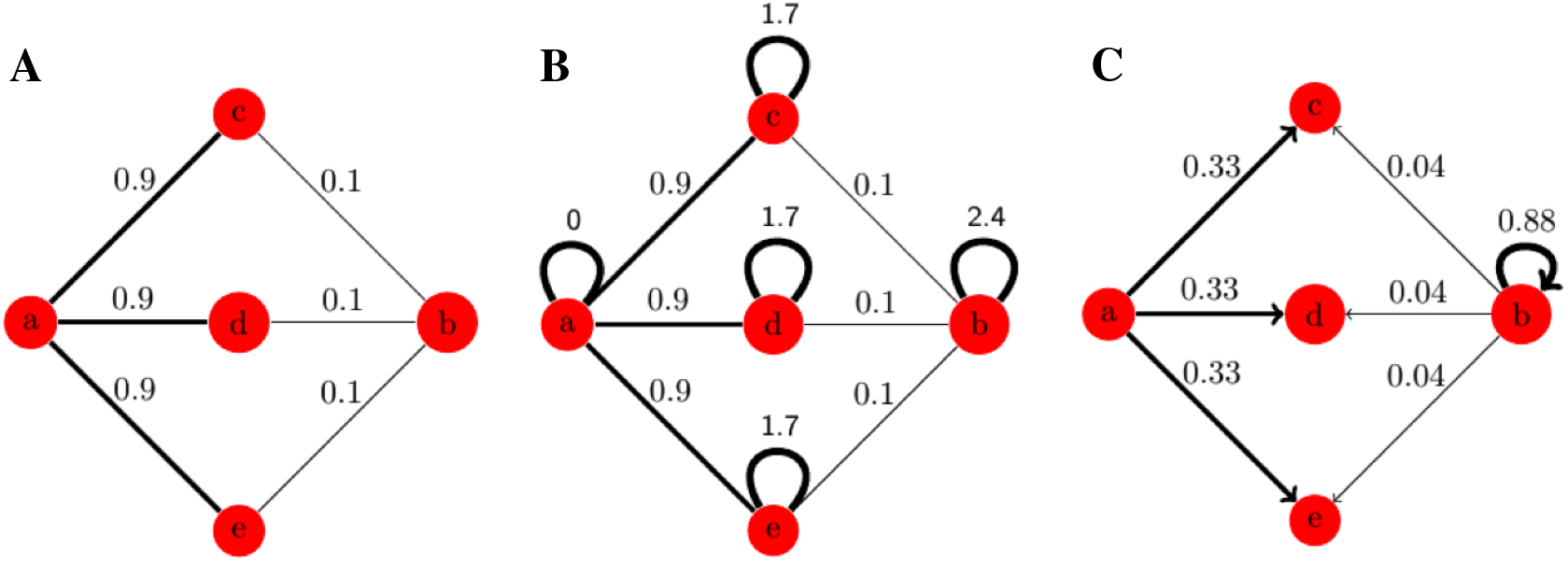
Edge strength normalization to maintain connectivity differences between a strongly connected and a weakly connected node. (A) A weighted graph with 5 nodes and 6 edges. (B) Adding self loops to nodes with weaker connections in order to normalize the probabilities. (C) Transition probabilities from nodes a and b after normalization (the transition probabilities from nodes c, d, and e are not included in this figure).

To overcome this problem, we add a self loop to nodes with weaker connections. To implement this, we find the node with maximum degree in the network. For every other node, we subtract the degree of that node from the maximum degree and add that as a self edge to the node. Therefore, the new degree matrix is *D′* = *d_max_I*, and the new adjacency matrix is 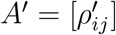, where:

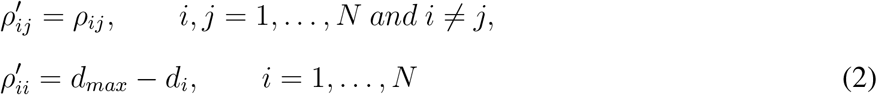

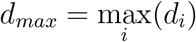, *i* = 1, …, *N* and *I* is the identity matrix.

### Hitting Time

We run random walk models on the graph with the new transition probability matrix 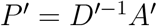 to calculate the hitting time matrix *H* = [*h_ij_*]. *h_ij_*, the hitting time from node *i* to node *j* is the expected number of hops to visit node *j* for the first time, for a random walk started at node *i*.

Hitting time is an asymmetric measure, meaning that *h_ij_* and *h_ji_* might be different. For example for a lollipop graph, the hitting times from the nodes on the complete component to nodes on the chain are much larger than the reverse direction, because a random walker spends more time in the complete component. We compute the hitting times between pairs of nodes using the graph-Laplacian method introduced in spectral graph theory Aldous and Fill (2002). This method is advantageous as it does not require the exact knowledge of the adjacency matrix, instead using a probabilistic approximation of the adjacency matrix of the network. Following Lovász and Simonovits (1993), we calculated the normalized graph Laplacian as:

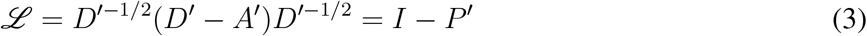

where, *D′* is the degree matrix and *A′* is the adjacency matrix of the graph after normalization as defined in the main text. We used the eigenvalues and eigenvectors of 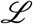 to calculate the hitting time matrix *H* = [*h_ij_*] Lovász and Simonovits (1993);

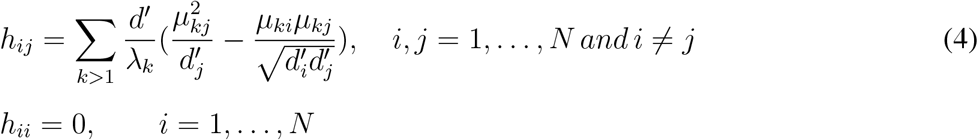

where, 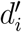 is the degree of node *i, i* = 1, …, *N*, and *d′* is the sum of all degrees (after normalization, see main text). 0 = λ_1_ < λ_2_ < · · · < *λ_n_* are the *n* eigenvalues of 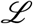, and *μ_kj_* is the *j^th^* element of *k^th^* eigenvector of 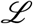 Lovász and Simonovits (1993).

Adding the self loops in the normalization step does not make the graphs reducible or periodic, meeting the requirements of the hitting time calculation we use here Norris (1997). Code for analysis in this project can be found under the first author’s name on github: https://github.com/SNaGLab.

## RESULTS

To detect and characterize linear chains of nodes, we focus on the random-walk measure of connectivity hitting time Lovász and Simonovits (1993) defined above. In synthetic graphs and estimated networks, a node is a point in the graph (or area of the brain) and an edge is a connection between two nodes. Hitting time between nodes *i* and *j* is a random variable describing the number of steps to get from node *i* to node *j* for the first time (represented as *h_ij_*) during a random walk, a measure equivalent to mean first-passage time Avena-Koenigsberger et al. (2017). Diffusion measures of networks, like hitting time, are becoming more commonly used and are the focus of active research Goñi et al. (2013); Lambiotte, Delvenne, and Barahona (2014); Shen and Meyer (2008). Diffusion-based measures carry significant methodological advantages. First, they overcome common issues caused when thresholding is used to define binary connections Goulas, Schaefer, and Margulies (2015); Reijneveld, Ponten, Berendse, and Stam (2007); Rubinov and Sporns (2011); Zalesky, Fornito, and Bullmore (2010). Second, they do not require perfect knowledge of the network to make a robust estimation of connectivity (i.e. it does not require the exact adjacency matrix Lovász and Simonovits (1993)). Third, measures like hitting time are asymmetric, meaning hitting time from one node to another may be different from the return trip, giving the best opportunity to identify extremeness in connectivity Lovász and Simonovits (1993). Here, we will use “hitting time” to refer to the expected number of edges to be traversed rather than the variable itself and “hitting-time distribution” to be the subject-average distribution of the expected number of edges to be traversed when moving between combinations of nodes. We begin by looking at the relationship between extreme hitting times and synthetic graph structure and then extend those findings to a publicly available functional magnetic resonance imaging (fMRI) dataset.

### How does the relative isolation of a linear chain of nodes change the distribution of connectivity in a synthetic network?

We consider a chain of sequentially connected nodes as a model for a hierarchical processing stream. Formally, a chain of sequentially connected nodes can be described as *N* nodes arranged in a line, so that there is an edge between nodes *i* and *i* + 1 for *i* = 1, …, *N* − 1, and no edges between nodes *i* and *j* where, *j* ≠ *i* − 1, *i* + 1. Theoretical results have found that a chain of sequentially-connected nodes attached to a fully-connected network (i.e. a lollipop graph) results in maximal hitting times when the chain is a third of the network Brightwell and Winkler (1990). We now compare the distribution of hitting times over nodes in a lollipop graph to small-world Watts and Strogatz (1998), random (Erdoõs-Rényi) Erdoõs and Rényi (1959), and complete synthetic graphs in Fig. 2. Each graph consists of 100 nodes. Random and small-world graphs are an average of 100 configurations. Linear chains of nodes result in larger hitting times which produce increased skewness in the hitting-time distribution. For example, in Fig. 2, panels A and D represent two extreme examples of hitting-time distribution in a network. Due to the presence of a path in A, the hitting time has a long tail (non-zero probability) that extends to large values (above 120,000). While in D each node is fully connected to every other node and hence, hitting time is the same across all pairs (in this case, 100) and the distribution is a single value with no tail. We focus on Kelley skewness Kelley (1923), as our measure of skewness because it directly compares the tails of the distribution. Kelley skewness (hereafter just skewness) therefore provides a more robust separation of extreme cases from changes in the interior of the distribution.

**Figure 2:**
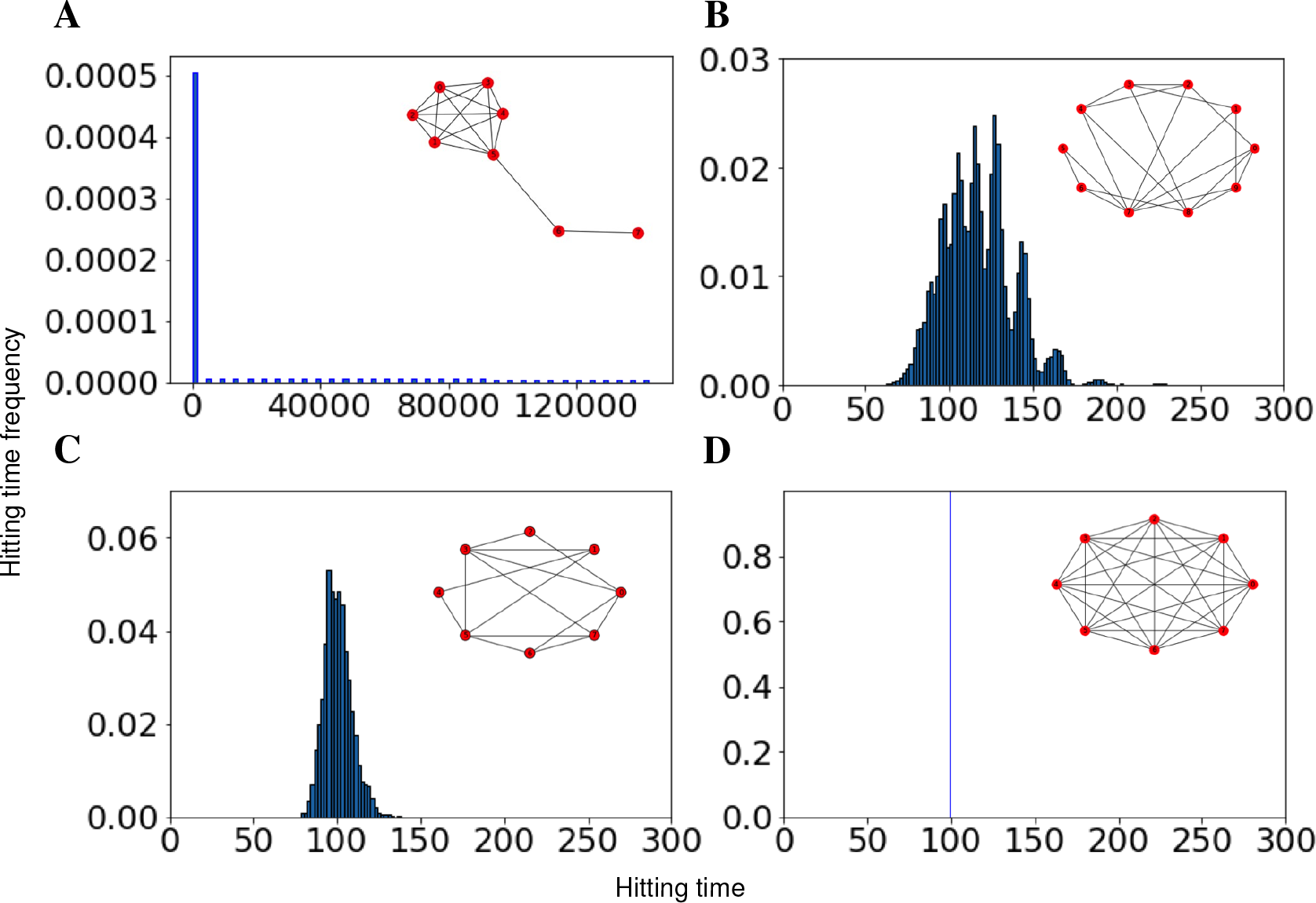
Hierarchical processing streams in lollipop graphs produce extremely long hitting times. Hitting-time distributions for (A) lollipop, (B) small world, (C) random and (D) complete graphs with 100 nodes (averaged over 100 runs for small world and random networks). The graphs on top of the distributions are smaller representations of the graphs used to generate the distributions. Axis scales change significantly with graph type (e.g. lollipop hitting time is several orders of magnitude larger than the other graphs).

Although perfectly isolated linear chains of nodes produce extreme hitting times, it is possible that even weak connections to the chain might significantly reduce hitting times to nodes on the chain. To characterize changes in the hitting-time distribution when a linear chain of nodes is not perfectly isolated, we begin with a random graph and alter its connectivity to isolate a linear chain of nodes (Fig. 3). Beginning with 50 nodes, edges of weight 1 were added between each pair of nodes with a probability of 0.6. We then randomly chose 10 connected nodes in the graph (1/5 of the graph) and reduced the weight of edges between those connected nodes and the rest of the network by 0.05 for 19 iterations. This process created a linear subgraph that becomes progressively more isolated until it resembles the stick of a lollipop graph. As a control for reduced connectivity across the network as a whole, we took the same graph we started with above and reduced all existing edge weights by 0.05 for 19 iterations. To preserve the reduction in overall connectivity (edge weights are typically normalized by the total connectivity of a node) edge weight reductions were added back as self loops (see Materials and Methods). These self loops are required as part of the random-walk to preserve the physiological principle that reduced neuronal activity would result in a reduction of connectivity (not just a shift between connections).

**Figure 3:**
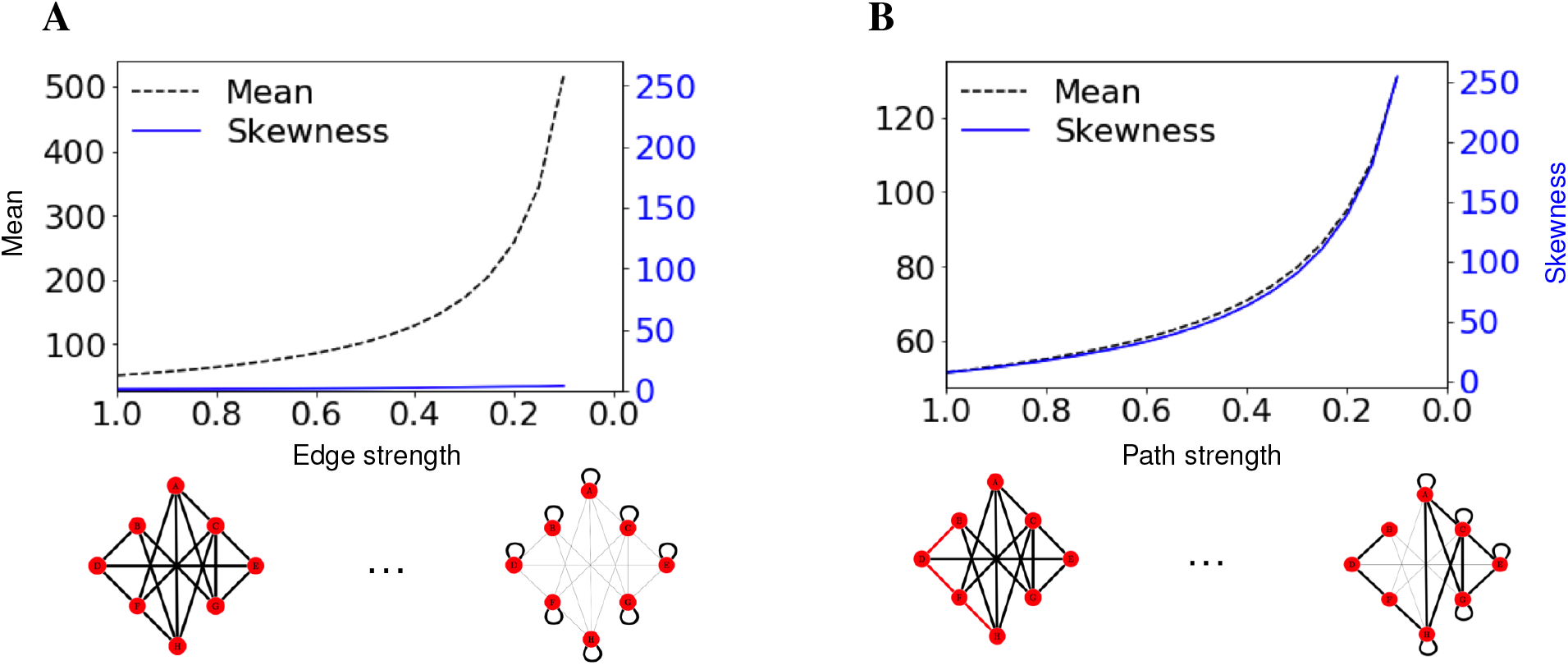
The skewness of the hitting-time distribution distinguishes a reduction of overall connectivity from a subgraph that becomes more linear. (A) Mean and skewness of hitting-time distribution as the strength of all connections is reduced by 0.05 for 19 iterations. Toy networks for this transition are represented on the x-axis. Reductions in connectivity are added as self loops. (B) Mean and skewness of hitting-time distribution as the strength of connections between linear component and the rest of the graph is reduced by 0.05 for 19 iterations. Toy networks on the x-axis represent synthetic graphs with a linear subgraph as the path (red) is made more linear.

Average hitting time increases as the chain of nodes becomes more isolated but also when the graph becomes more disconnected as a whole. Hence, mean hitting time does not distinguish between these two scenarios. However, in our simulations, skewness changed significantly as the linear subgraph becomes more isolated but only minimally when the average connectivity of the whole graph decreased. Skewness also increased with the relative isolation of the chain of nodes but was present even when each node in the chain was somewhat connected to the rest of the graph (Fig. 3).

We have shown that a lollipop component present in a graph results in significant increase of (Kelley) skewness of hitting-time distribution but is this relationship true in heterogeneous topologies? It is important to note that other changes in graph structure may also result in extreme hitting times. One commonly employed graph measure is modularity, the extent to which the graph can be easily separated into different communities.

To evaluate the effect of modularity on hitting-time distribution, we have tested a large number of networks with different levels of Louvain modularity and numbers of chain motifs (3 node linear components). We generated random networks each with 100 nodes varying the number of edges. To allow a comparison across a given number of edges, we generated multiple graphs with the same number of edges by choosing *k* edges uniformly from the full possible set of edges. *k* was varied from 200 to 1000 in intervals of 50 and 1000 to 2500 in intervals of 100. The range of the average degree is [4, 50], with a range of Louvain modularity of [.1,.6]. Keeping only those graphs that were connected resulted in 15,243 graphs for comparison. Using a linear model, we sought to explain skewness as a function of modularity, number of edges, and number of chain motifs. Number of edges (*p* < .001, *t*(15241) = 4.16, *β* = −.0052), modularity (*p* < .001, *t*(15241) = 37.4, *β* = 239), and the number of chain motifs (*p* < .001, *t*(15241) = 13.8, *β* = 9.5) all independently explain some variance in skewness. In the remaining analysis of brain networks, we therefore test whether or not nodes with extreme hitting times also become less chain like.

### Are there subnetworks in resting-state cortex that have properties similar to a linear chain of nodes?

Motivated by the above simulations, we utilized the skewness of the hitting-time distribution to identify potential linear chains of nodes in cortical connectivity data. The brain is made up of a large number of highly interconnected regions Cherniak (1990) evolved to efficiently integrate a variety of sources of information Friston (2010) that can be represented as a network. Graph-theoretic models of the brain have been used to effectively segment commonly associated regions of the brain into large-scale networks and describe the properties of brain information processing (for review see E. Bullmore and Sporns (2009)) in health and disorder Bassett and Sporns (2017); Fox and Greicius (2010). Characterizations of brain network changes in development or psychiatric disorder often utilize graph measures like efficiency Latora and Marchiori (2001) and small-worldness Watts and Strogatz (1998), which typically include the average path length in their definition (for common measurement descriptions, see Achard and Bullmore (2007)). Even measures that may not directly utilize the average path length (e.g. modularity Newman and Girvan (2004); Stiso and Bassett (2018)) sometimes rely on community detection methods that incorporate the average path length. The use of an average path-length rests on the assumption that path lengths in that network are normally distributed and so can lead to the mischaracterization of the topology of the network. The concern arises because of the use of an average and is present in both traditional and diffusion-based graph measures. Overcoming this assumption requires the use of specific subnetwork models (for example, see Khambhati, Medaglia, Karuza, Thompson-Schill, and Bassett (2018)) or the capture of deviations from normality in the path-length distribution. Here, we use Kelley skewness of the hitting-time distribution to distinguish changes in the network as a whole from the presence of network topologies resembling hierarchical processing streams.

To test for the presence of skewness in cortical connectivity, we generated a hitting-time measure of connectivity (see materials and methods) for resting-state functional data from neurotypical participants who were part of a large open-source dataset (LA5c, UCLA Consortium for Neuropsychiatric Phenomics Poldrack et al. (2016), see supporting information). Network nodes were 180 anatomical regions from the multi-modal parcellation of Glasser et al. Glasser et al. (2016). The average hitting-time distribution of neurotypical resting-state functional connectivity is positively skewed (Kelley skewness of 15.04 and Pearson’s coefficient of skewness of 2.3), Fig.4A. A D’Agostino-Pearson test Trujillo-Ortiz and Hernandez-Walls (2003) showed that as a whole the hitting times were not normally distributed (*Z*(*skew*) = 110, 3496, *χ*^2^(2) = 17864.8071, *p* < 0.001).

**Figure 4:**
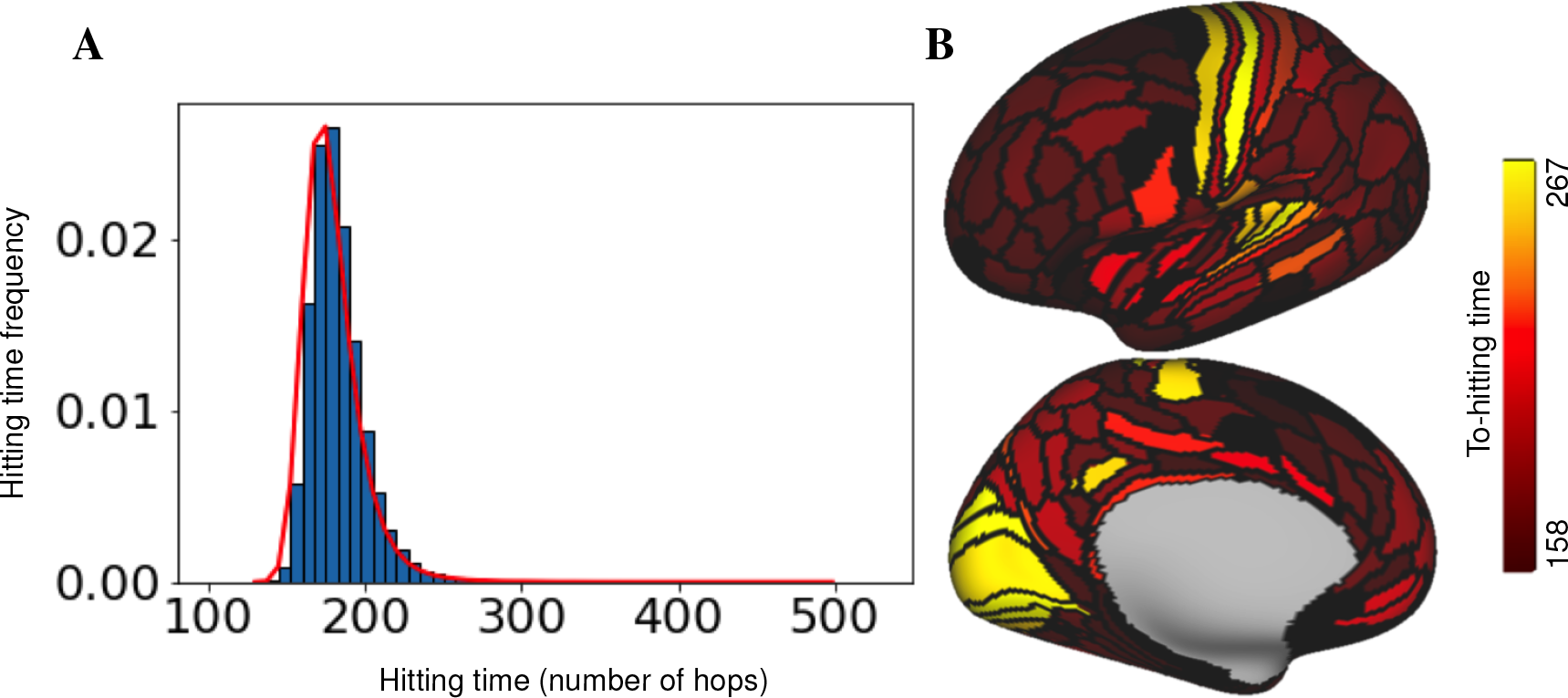
Hitting-time measures of resting-state functional connectivity in neurotypical participants are positively skewed. (A) Average normalized hitting-time distribution for control subjects during resting-state from the publicly available LA5c study. (B) Average to-hitting time from all other regions of cortex for lateral (top) and medial (bottom) left maps thresholded to [158(10%), 267(95%)]. The range of data is [129, 310]. Primary auditory, visual, and somatosensory cortices have the largest to-hitting times in the cortex.

Although skewness of a distribution can originate from many sources, ranging from a smooth shift of the distribution as a whole to far ranging outliers, the particular skewness measure used here (Kelley skewness) directly compares the extremes of the distribution (90% compared to 10%), limiting the potential causes of the skewness. Limiting our test for skewness to the tails of the distribution is consistent with our aim of identifying changes in linear-chain topologies, which have been shown to produce maximal hitting times in lollipop networks (see above). The primary auditory, visual, and somatosensory hierarchies, show the largest average to-hitting times (Fig. 4B), and are therefore possibly related to chain-like network topologies. It is important to note that even the use of a Kelley skewness metric does not guarantee the presence of chain-like network topologies. In fact, random graphs generated from a stochastic block model that precluded chain-like topologies exhibited Kelley skewness explained by modularity and node degree. We generated 100 graphs with 180 nodes from a stochastic block model. To define groups and mixing structure, we fixed the probability of connections within communities to be (*p* = 0.7) and between communities to be (*q* = 0.1). To expand the range of possible modularity, we randomly picked the number of nodes in each community until we reach 180 (if the total number of nodes passes 180, we reduce the size of the last community to have a total of 180 nodes in the graph). After removing the null values, we ended up with 89 graphs with number of communities from 2 to 6 and Louvain modularity levels of 0.03 to 0.42. These SBM models contain Kelley skewness which is explained by both modularity (*β* = −1.591*e* + 03, *t*(86) = −10.76, *p* < 0.001) and degree (*β* = −8.617*e* − 02, *t*(86) = −7.39, *p* < 0.001). Because Kelley skewness may arise from multiple sources, we next look for changes in the skewness of the hitting-time distribution during a task and ask whether those changes are related to regions of the brain with the largest hitting times during resting scans, and then, whether those areas with the largest rest hitting times also become less chain-like.

### How are linear-chain subnetworks changed by task demands?

To better interpret the skewness of cortical networks during resting-state fMRI, we sought to test whether hitting times become more or less skewed during task performance and which connectivity changes underlie those shifts. We compared resting-state and balloon analogue risk task (BART) functional connectivity from the LA5c study. The BART is a paradigm designed to study risk taking in an experimental setting. Participants in the BART decide whether or not to pump a balloon that is at risk of popping. The BART is defined by visual input and motor responses without structured auditory stimulation.

Hitting times between cortical areas were calculated for fMRI data collected during the performance of the BART task using the same processing pipeline as for the resting-state scans. To test for differences in skewness, we then ran a linear mixed effect model (lme in R) of skewness of the hitting-time distributions (dependent variable) modeling task (with resting-state as a reference), gender, and age as independent variables. The task variable was treated as a random effect (BART and resting-state points were paired by participant), which characterizes idiosyncratic variation that is due to individual differences. In our first model, we found significant difference in skewness for control subjects between BART and rest (*β* = −10.25, *t*(118) = 0.79, *p* < 0.001), see Fig. 5. The skewness of hitting-time distribution is significantly reduced in the BART (*μ* = 5.48, *σ* = 3.96) compared to rest (*μ* = 15.73, *σ* = 8.49). Age and gender did not significantly explain variance in this model. We next sought to test whether this skewness could be related to nodes with extreme hitting times and whether those nodes define network topologies that become less chain-like.

**Figure 5:**
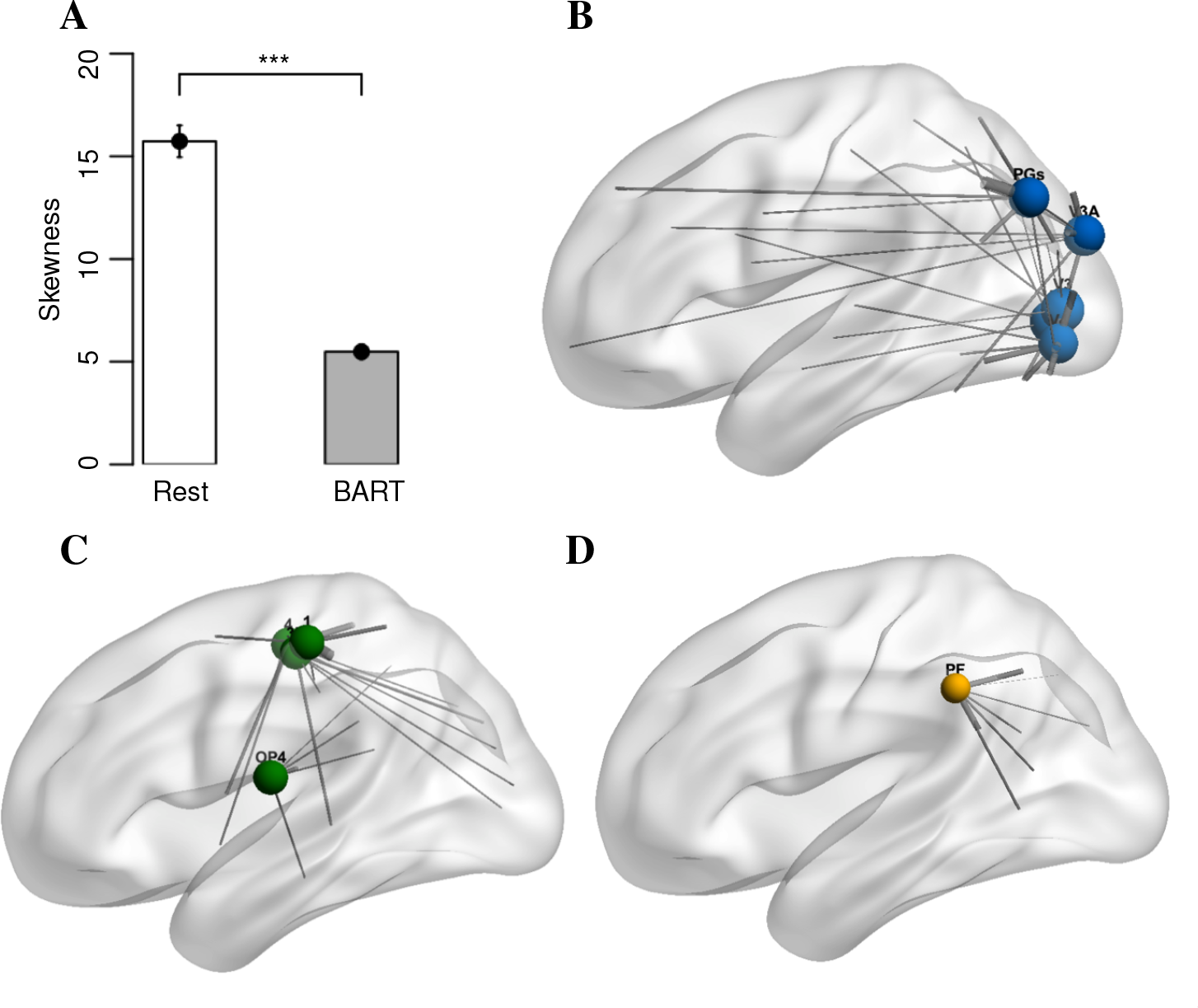
The distribution of skewness vs. task for control subjects. p-values significance codes:. = 0.1, ∗ =*<* 0.05, ∗∗ =*<* 0.01, ∗ ∗ ∗ =*<* 0.001. The skewness of hitting-time distribution for control subjects is significantly smaller when the subjects are engaged in BART task compared to rest. The 10 nodes with largest hitting time changes are (B) V2, V3, V4, V3A and PGs (visual), (C) 1, 4, 3b and OP4 (motor) and (D) PF. The size of each node represents the magnitude of difference of average to-hitting times (range from 19 to 30.2) and thickness of each edge represents the magnitude of difference of partial correlation in BART compared to rest.

To identify those nodes related to differences between rest and task, we first ask which nodes had the largest hitting-time changes. The ten regions with the largest hitting-time changes (paired t-tests comparing task and rest hitting times, significantly different with *p* < 0.05, Bonferroni corrected) are ‘V2’, ‘V3’, ‘V4’, ‘V3A’ and ‘PGs’ within the visual cortex, ‘1’, ‘4’, ‘3b’ and ‘OP4’ within somatosensory cortex, and the area ‘PF’, Fig. 5B-D. Regions are labeled according to Glasser et al. (2016). These nodes, which show decreased hitting times during task performance, overlap heavily with the visual and motor processing streams and correspond to many of the nodes with the largest hitting times during rest scans. This reduction in hitting time in the visual and motor pathways during the BART provides support for the role of these pathways in skewness but does not address whether differences in the chain-like topology is responsible for the change in hitting times.

To add support for the role of chain-like topologies in large hitting times in brain data, we calculated a chain-index for each node that provides a measure of how similar to an isolated chain the local connectivity of the node is. For every node in the network located on a three node chain motif, we define a chain index by focusing on its two strongest connections. A node is located on a chain motif if its neighbors with the two strongest connections are stronger than the remaining connections and if the two strongest connections have a significantly weaker connection with each other. Assuming that node *i* is on a chain motif and has *N_i_* neighbors and nodes *i*_1_ and *i*_2_ have the strongest connections to node *i*, we define chain index for node i as:

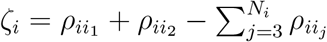

If a node is located on a perfect chain, the chain index will be at its maximum. If a node has equal connections to every other node in the network (the least chain-like network), the chain index will be a large negative number that depends on the number of nodes in the network.

Comparing connectivity during the BART to rest, the largest reductions in the chain index were for a set of nodes that overlapped heavily with those nodes with the largest hitting time (8 of the 10 nodes described above), including ‘V2’, ‘V3’, ‘V3A’, ‘V6’ and ‘V6A’ within the visual cortex,’1’,’4’,‘3b’ and ‘OP4’ within somatosensory cortex, and the area ‘PF’. Labels are according to Glasser et al. (2016). In accordance, the nodes with the largest hitting-time changes also show increased connectivity during the BART. Connections from nodes with the largest hitting times that have changed significantly (paired t-tests comparing connections from each node during task and rest, *p* < 0.05 Bonferroni corrected) are indicated in Fig. 5 by gray lines, with the thickness of the line indicating the size of the change. During task performance, nodes in task-related sensory streams (which have large hitting times during rest) have smaller hitting times, becoming less like isolated chains and instead more widely integrated.

### Does the characterization of network topology have value in understanding and diagnosing psychiatric disorders?

Brain network efficiency, which is commonly defined by the mean distance between nodes, has been shown to change in disorders such as Alzheimer’s Dennis and Thompson (2014), schizophrenia Besnard et al. (2018); Li et al. (2017), and others Cheng et al. (2016). Reductions in measures of efficiency that utilize mean distance could be due to: 1. Reduced overall connectivity; or, 2. Subnetwork changes that lead to skewed hitting-time distributions. As described in Fig. 3, we can distinguish these possibilities by focusing on skewness. If skewness changes, the differences between psychiatric populations and controls is more likely to be due to subnetwork changes than a change in over all connectivity. We ran an ordinary least squares regression model with skewness of the resting-state hitting-time distribution as the dependent variable, and group (dummy coded, reference controls), gender (dummy coded, reference females), and age (mean centered, linear) as independent variables. We analyzed resting-state functional data from four patient groups, control, schizophrenia, bipolar and attention deficit hyperactivity disorder (ADHD). We found significant differences in skewness between schizophrenia and control populations (*β* = −5.130, *t*(252) = 1.268, *p* < 0.001), bipolar and control populations (*β* = −4.060, *t*(252) = 1.324, *p* < 0.001), and a trend-level difference between ADHD patients and control populations (*β* = −2.445, *t*(252) = 1.324, *p* = 0.066), see Fig. 6. Gender and age did not significantly explain variance in this model.

**Figure 6:**
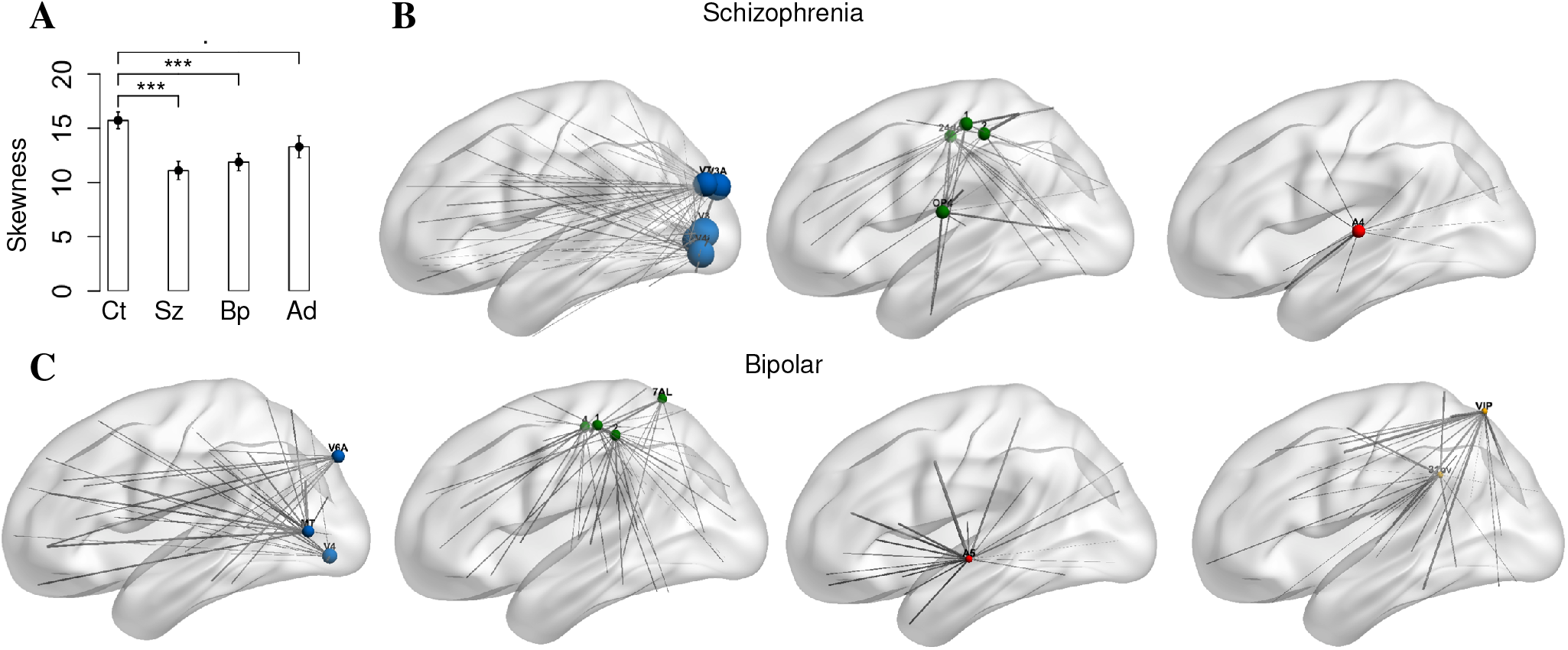
Skewness of the hitting-time distribution is significantly different across patient groups. (A) Distribution of skewness of the hitting-time distributions in patient and control groups during resting-state scans. Ct, Sz, Bp and Ad stand for control, schizophrenia, bipolar and ADHD, respectively. Significance codes:. = 0.1, * =< 0.05, ** =< 0.01, * * * =< 0.001. The skewness of hitting-time distribution is significantly smaller for subjects with schizophrenia and bipolar disorders compared to neurotypicals. The 10 nodes with the largest change in hitting time for subjects with schizophrenia include V2, V3, V4, V3A and V7 (visual, top blue), 1, 2, 24dd and 4 (motor, top green) and A4 (auditory, top red). The 10 nodes with largest change of hitting time for subjects with bipolar include MT, V4, V3A and V6A (visual, bottom blue), 1, 2, 4 and 7AL (motor, bottom green), A5 (auditory, bottom red) and VIP and 31pv (bottom yellow). The size of each node represents the magnitude of difference of average to-hitting times (range from 7.1 to 33.4) and thickness of each edge represents the magnitude of difference of partial correlation in psychiatric disorder compared to control subjects during rest.

To identify those nodes related to skewness differences between schizophrenia, bipolar disorder, and controls, we first ask which nodes had the largest hitting-time changes. The ten regions with the largest hitting-time changes between (t-tests comparing schizophrenia and control groups for each region, significantly different with *p* < 0.05 Bonferroni corrected) were ‘V2’, ‘V3’, ‘V4’, ‘V3A’ and ‘V7’ within the visual cortex, ‘1’, ‘2’, ‘4’ and ‘24dd’ within somatosensory cortex and ‘A4’ within the auditory cortex for subjects with schizophrenia. For subjects with bipolar disorder, the ten regions with the largest hitting-time changes (significantly different in t-tests comparing bipolar and control groups with *p* < 0.05 Bonferroni corrected) were ‘MT’, ‘V4’, ‘V3A’ and ‘V6A’ within visual cortex, ‘1’, ‘2’, ‘4’ and ‘7AL’ within somatosensory cortex, ‘A5’ within auditory cortex and ‘VIP’ and ‘31pv’. Regions are labeled according to Glasser et al. (2016). Connections that have changed significantly (t-tests comparing connections from each node between groups, *p* < 0.05 Bonferroni corrected) from these nodes are indicated by gray lines with the thickness of the line indicating the size of the change.

We find evidence that individuals with schizophrenia and bipolar disorder have less skewed hitting-time distributions than controls during resting-state fMRI. The regions of cortex with the largest hitting-time reductions between patient and control populations are in sensory/motor cortex and overlap with many of the same regions that have extremely large to-hitting times during rest in controls. It is possible that the large hitting-time values found in these regions of the cortex are related to the theoretical finding that linear-chains of nodes produce maximal hitting times and that a reduction in hitting time in these regions occurs when the nodes are part of a topology that is less chain like. We therefore also compared the chain index for schizophrenia, bipolar and control groups. The ten regions with the largest changes of chain index (t-tests comparing schizophrenia and control groups for each region, significantly different with *p* < 0.05 Bonferroni corrected) are ‘V2’, ‘V3’, ‘V4’, ‘V3A’ and ‘V7’ within the visual cortex, ‘1’, ‘OP4’ and ‘24dd’ within somatosensory cortex and ‘A4’ and ‘A5’ within the auditory cortex for subjects with schizophrenia. For subjects with bipolar disorder, the ten regions with the largest changes of chain index (significantly different in t-tests comparing bipolar and control groups with *p* < 0.05 Bonferroni corrected) are ‘V4’, ‘V3A’ and ‘V6A’ within visual cortex, ‘1’, ‘2’, ‘4’ and ‘7AL’ within somatosensory cortex, ‘A5’ within auditory cortex and ‘VIP’ and ‘PFcm’. The convergence of evidence of changes in extreme hitting-time values in sensory areas of cortex and those same areas being connected in a less chain-like topology in schizophrenia and bipolar disorder is consistent with our hypothesis that path-length changes in these populations are likely to be related to sub-network topology shifts and not changes the network on average.

## DISCUSSION

We have presented evidence that the skewness of connectivity in cortical brain networks can be used to infer likely network topology changes that improve our understanding of information processing in the brain. Using random graphs we showed that the isolation of linear motifs is one prominent cause of skewness in hitting time even in the presence of mixed topologies. We then showed that skewness, but not average of brain-connectivity distributions is related to psychiatric diagnosis. We confirmed that these differences in skewness were related to a linear-chain topology by testing for changes in a chain index. In networks for which connectivity is positively skewed, a change in subnetwork topology (and possibly a different brain state) is a more parsimonious explanation than changes in average connectivity. These topology changes are focused in sensory areas of the brain and when compared to changes brought about during task performance, provide an initial mechanistic link between resting-state connectivity changes and clinical diagnosis.

Extremely large hitting times can be linked to linear chain topologies through theoretical work showing that lollipop networks result in maximal hitting times Brightwell and Winkler (1990). We have shown using a toy problem that the extremeness of hitting-time values scales with how isolated a linear chain of nodes is and that the presence of chain motifs is related to extreme hitting-time values even in random networks with mixed topologies above and beyond modularity of the network. Resting brain networks also have extreme hitting times that are likely related to hierarchical processing in sensory cortex. When they are most isolated from the rest of the network, hierarchical processing streams, resemble the chain of nodes on the linear component of a lollipop graph. This parallel motivates the use of extremely long random walks between brain areas as a measure of the presence, and relative isolation, of hierarchical processing streams. In particular, hitting time of a region of interest can be utilized to detect presence of linear components likely to be hierarchical processing streams. A central dogma of neuroscience is that sensory representations are constructed hierarchically Hubel and Wiese (1962); Kikuchi, Horwitz, and Mishkin (2010); Van Essen and Maunsell (1983). Hierarchical processing streams have also been a focal component of computer vision models since 1971 Giebel (1971) and a significant contributor to the success of modern convolutional neural networks LeCun, Bengio, and Hinton (2015). Their foundational nature has made the study of hierarchical processing streams the focus of targeted analyses (e.g. Sepulcre, Sabuncu, Yeo, Liu, and Johnson (2012)). Here, we showed that nodes from sensory and motor areas of the brain have extreme hitting times which contribute to Kelley skewness. By comparing resting to task-based network topologies, we can show that decreases in hitting time are also associated with sensory hierarchies becoming less chain like.

Task changes in functional connectivity can be observed as changes in functional brain network topology. Previous work has highlighted the central role of path length and the integration of isolated paths in function. In Goni et al. Goñi et al. (2014), a notion of path transitivity – which accounted not only for the shortest path, but also local detours along that path – was the best predictor of functional connectivity. Similar notions of distributed communicability (related to the diffusion of information over the network) were used to quantify the disruption of the global communication in the cortex that was triggered by the pharmacogenetic inactivation of the amygdala Grayson et al. (2016), and to detect changes in functional connectivity after a stroke Crofts et al. (2011). A thorough review of these concepts and their relationships to the notion of mean first-passage time, which is equivalent to the mean hitting time, is provided in Avena-Koenigsberger et al. Avena-Koenigsberger et al. (2017). Because the mean hitting time conflates overall changes in connectivity with changes to a subnetwork, these changes may be better explained using Kelley skewness, which specifically focuses on extreme values and so provides a mechanism to identify potential subnetwork changes. Extreme values contributing to connectivity skewness were associated with sensory areas of the brain specifically associated with the task. Hitting times in these sensory areas of the brain became shorter and less extreme during BART task performance. Sensory areas became more strongly connected to distant areas throughout the brain. The nodes with the largest reductions in hitting time were found in brain sensory areas related to the BART task (somatosensory and visual areas). One additional area also showed a decrease in hitting time, PF. PF is located in the inferior parietal lobule and is thought to be related to risk processing Weber and Huettel (2008). In line with its role in the processing of visual magnitude, it showed increased connectivity with visual inputs. Broadly, the introduction of a task caused sensory processing streams to become better connected to other task relevant areas, and less chain-like. This result is somewhat counterintuitive since hierarchical processing pipelines are often described as most distinct when active. We do find evidence of increased connectivity within sensory networks but these increases in strength within the sensory processing stream are offset by wider integration making their network topology less chain like. In addition, whether the processing pipeline becomes more or less integrated depends on the calculation underlying the transition probability between areas. When similar models are constructed using the raw correlation values, group and task differences in the same data set are consistent but in the opposite direction Rezaeinia and Carter (2017). This is likely due to the redundant connections and task event correlations Cole et al. (2018) included in raw correlation models. Here, we focused on the partial correlation of brain-region time-series which minimizes redundant connections, following the state-of-the-art in the field Smith et al. (2011). In spite of the complexities raised, the distribution of task relevant information throughout the brain is consistent with what would be most likely to improve BART task performance and supports the interpretation of sensory hierarchies as existing as relatively isolated linear networks topologies during rest. In future work it would be helpful to incorporate additional multifaceted tasks to generalize these findings to the incorporation of sensory information under other constraints.

Skewness of the connectivity distribution also explains cortical-network differences between psychiatric diagnoses. Resting functional connectivity in a large neurotypical population is significantly positively skewed. In such a case, average efficiency for such a network would be biased and less representative of the network as a whole. An important finding is that changes in connectivity between clinical and control populations are due to changes in skewness rather than average differences. In fact, the median of connectivity measures changed in the opposite direction with respect to the average. Network connectivity changes between clinical and control populations are therefore due to a subset of connections rather than the network as a whole. The identification of specific cortical regions involved in topological changes between neurotypical and clinical populations provides an opportunity to better understand functional changes that occur in those populations as well as opportunities for improving diagnosis. Task performance reduced hitting-time skewness by increasing the connectivity between sensory areas and the rest of the cortical network. Qualitatively similar changes are seen in clinical populations, implying further work exploring network topology changes to specific tasks may help characterize the atypical resting connectivity for individuals with a schizophrenia or bipolar diagnosis. A testable prediction from this implied mechanistic difference would be that individuals with these diagnoses spend less time in activities typically associated with resting fMRI (e.g. future planning).

The results presented were based on theoretical predictions and applied to a publicly available dataset in a rigorous manner. We would like to document the following caveats and qualifications. First, although linear components in networks produce maximal hitting times in theory, it is possible that other network topologies could also produce some degree of skewness. To answer this concern, we showed that in toy examples skewness was related to short linear graphs or linear graphs that were still connected to the rest of the graph. We also found that the hitting-time distribution of both small-world, lollipop, and random graphs are skewed but that skewness in both cases is dominated by linear paths, even with a linear-path that comprises significantly less than a third of the graph. It is, however, important to note that the number of linear topologies does not explain all of the variance in Kelley skewness and so there could be other contributing topologies, perhaps related to modularity and degree. In our human fMRI analyses, the network as a whole, did not become less connected (see supplementary materials) and those areas with extreme values become less chain-like when they have smaller hitting times. Thus, the relationship between extreme hitting times and linear paths is robust. In addition, past work on sensory hierarchies Hubel and Wiese (1962); Kikuchi et al. (2010); Van Essen and Maunsell (1983), recent work showing parallels between convolutional neural networks and sensory networks Güçlü and van Gerven (2015); Kell, Yamins, Shook, Norman-Haignere, and McDermott (2018); Khaligh-Razavi and Kriegeskorte (2014), and the integration of sensory networks during task performance (see above) are all consistent with the presence of linear components in the cortical network. Second, we focused on a cortical model of brain function and the absence of subcortical nodes could have affected the topology of the network model. However, the inclusion of subcortical connections to cortical network endpoints should not change connectivity measures since the path would still produce larger hitting times (the largest times may then be shifted to the middle of the sensory hierarchy).

## CONCLUSION

In conclusion, establishing a link between network topologies, hierarchical processing pipelines, task engagement, and psychiatric disorders provides an opportunity to interpret cortical network changes in the light of cognitive models of brain function. The interpretation of network connectivity and information processing topologies is an area of significant focus for neuroscience Fornito and Bullmore (2015). The widespread collection of rfMRI in particular provides a unique opportunity to extend this work to numerous psychiatric disorders and compare these findings with the growing body of open-source fMRI task data.

## SUPPORTIVE INFORMATION

### Data

We used the functional magnetic resonance imaging (fMRI) data from the LA5c Study Poldrack et al. (2016), collected by the UCLA Consortium for Neuropsychiatric Phenomics (CNP), which is funded by the NIH Roadmap Initiative. This data was obtained from the OpenfMRI database. Its accession number is ds000030. The dataset is formatted according to the Brain Imaging Data Structure K. J. Gorgolewski et al. (2016) (BIDS) standard. This study contains neuroimaging data from 290 participants. We ended up with a sample number of 255 subjects after removing subjects with missing functional measurements. In this sample, there are 119 healthy individuals (labeled as control), 49 individuals diagnosed with schizophrenia, 48 individuals diagnosed with bipolar disorder and lastly 39 individuals diagnosed with attention deficit hyperactivity disorder (ADHD). The focus of the LA5c Study is to understand memory and cognitive functional structures across patient groups. Therefore, the data set includes resting-state fMRI data as well as fMRI data collected during several different tasks. In this paper we focus on the analysis of resting-state and balloon analogue risk task (BART) fMRI data for the four specified groups. We focused on the BART because it has reliable visual input which has been shown to produce large-scale changes in sensory cortical hierarchies, which could be compared to the internally-driven resting-state network. In addition, we develop variants of the BART and so we have expertise with the task and any findings would have local applications. We obtained the preprocessed data which is de-identified, motion corrected and coregistered to Montreal Neurological Institute (MNI) standard space K. Gorgolewski, Durnez, and Poldrack (2017). We used FSL Jenkinson, Beckmann, Behrens, Woolrich, and Smith (2012); Smith et al. (2004) to correct for motion and apply a high-pass filter to remove low-frequency noise (cut-off frequency of 100 seconds). The data also includes potential confound regressors K. Gorgolewski et al. (2017). In order to remove the effect of motion artifacts, we used the 36-parameter motion regression technique introduced in Satter et al. Satterthwaite et al. (2013), which has been shown to be most effective in decoupling modular structure from subject motion Ciric et al. (2017).

### Parcellation

The fMRI data has the following parameters; the matrix consists of 64 × 64 voxels for 34 slices recorded with a TR of 2 seconds. In total, there are 147 time samples for resting-state data and 267 time samples for BART data. To extract a reliable cortical network for each participant, we reduced the number of nodes by averaging over voxels within an anatomical region. We used the multi-modal parcellation developed by Glasser, et al. Glasser et al. (2016) to map the fMRI data into a more sparse decomposition framework. There are 180 regions in each hemisphere, which increase the neuroanatomical precision for studying the structural and functional organization. This parcellation is based on multiple neurobiological properties, connectivity, functional and architecture, which improves the consistency across subjects.

## HITTING TIME VS. DEGREE DISTRIBUTION

To compare the ability of hitting time and degree distributions to distinguish the presence of linear components, we generated a random graph with *N* = 50 nodes in which a connection between each node randomly occurred with *p* = 0.6. We then attached a linear component of length 1 to the graph and kept increasing its length by 1 for 20 iterations (Fig. 7A). Skewness of the hitting-time distribution increases as we increase the length of linear component (Fig. 7B, blue/solid). The skewness of degree distribution remains close to constant (Fig. 7B, black/dashed). The change in other features of degree distribution such as mean or median do not reliably reflect the linear component either. Depending on the relative size of network and linear component, we might observe change in the degree distribution, but it is not consistent and is not most closely related to the presence of linear components.

**Figure 7:**
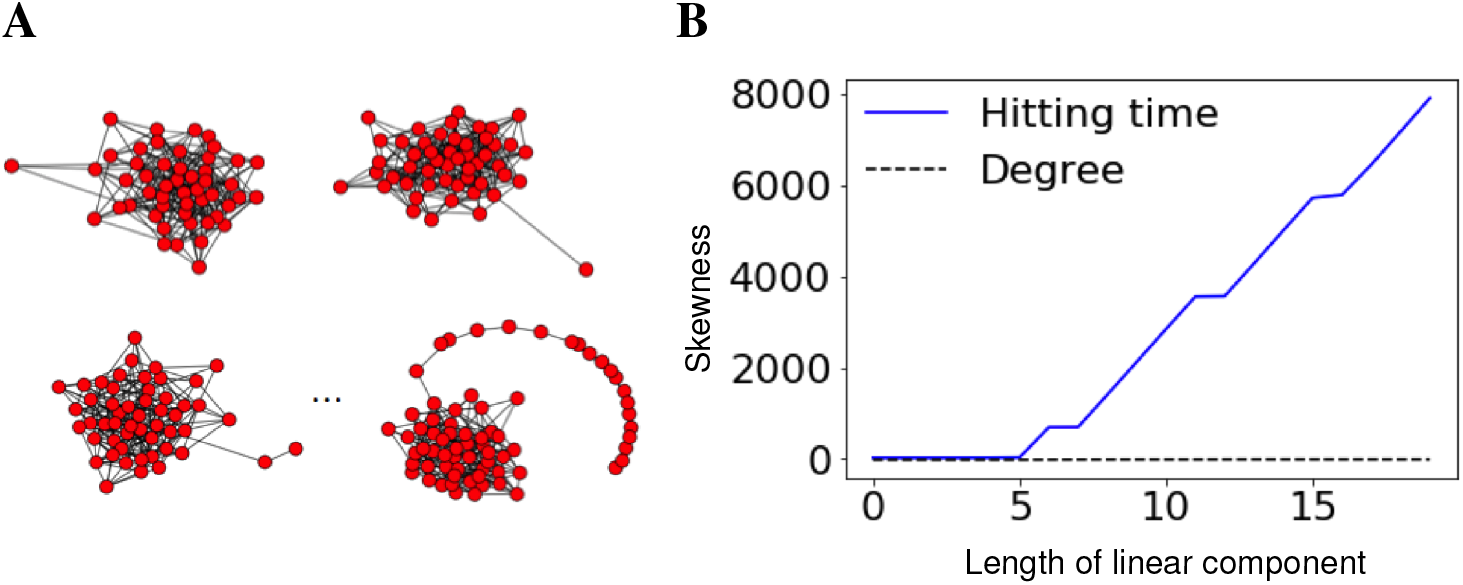
The hitting-time distribution of a graph becomes more skewed as the length of a linear subgraph increases. (A) We started with a random graph with 50 nodes, added a linear component of length 1 and increased the length of the linear component by 1 for 20 iterations. (B) Skewness of the hitting-time distribution increased significantly as the length of linear component increased. The degree distribution does not demonstrate a consistent relationship with linear path length.

## MEAN AND MEDIAN OF HITTING-TIME DISTRIBUTION

### Effect of task on mean and median of hitting-time distribution

Aligned with the tests for skewness, we seek the effect of task on mean and median of hitting-time distribution. We ran a linear mixed effect model by adding a random effect for participant (BART and resting-state points were paired by participant). We found that mean of hitting-time distribution is significantly smaller for BART compared to rest (*β* = −1.6, *t*(118) = 0.13, *p* < 0.001).

Finally, a linear mixed effects model on the median of hitting times reveals a significant positive effect of BART vs. rest, (*β* = 1.3, *t*(118) = 0.16, *p* < 0.001), Fig. 8.

**Figure 8:**
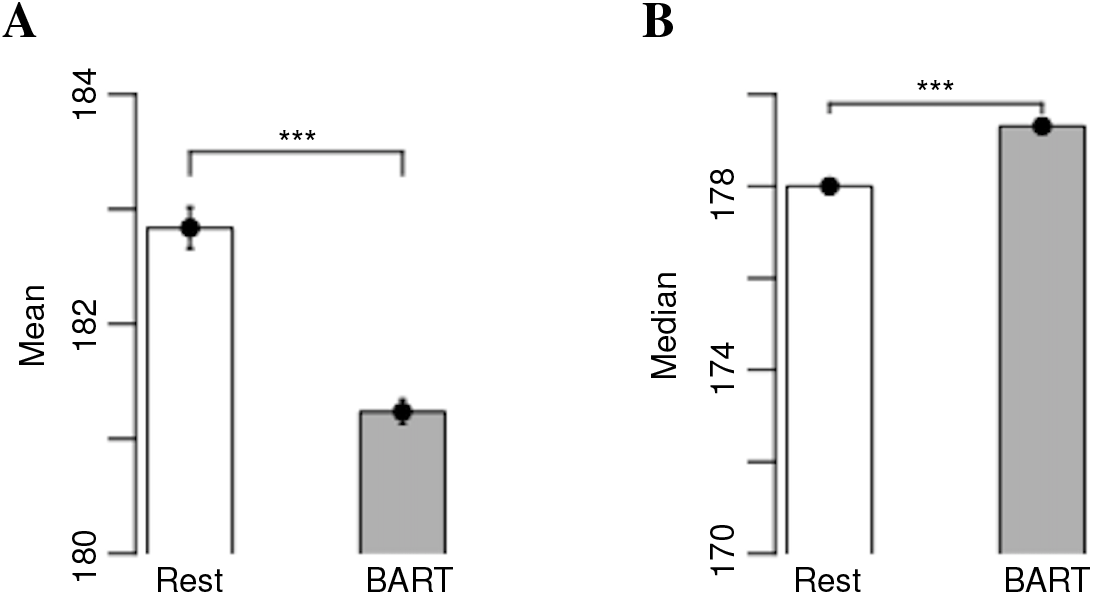
The distribution of (A) mean and (B) median vs. task for control subjects. p-values significance codes: = 0.1, * =< 0.05, ** =< 0.01, * * * =< 0.001.

### Effect of psychiatric disorders on mean and median of hitting-time distribution

To test the effect of psychiatric disorders on the average efficiency, we run an ordinary least squares regression model with mean of hitting-time distribution as the dependent variable. The independent variables are group (coded as a dummy variable with the control group as the reference group), gender (coded as a dummy variable with the female gender acted as the reference category) and age (mean centered, linear). We found significant differences between schizophrenia versus control (*β* = −1.113, *t*(252) = 0.255, *p* < 0.001), bipolar versus control (*β* = −1.064, *t*(252) = 0.252, *p* < 0.001) and ADHD patients versus control (*β* = −0.835, *t*(252) = 0.266, *p* = 0.002). The qualitative results are similar to skewness, but the coefficients are approximately one quarter the magnitude of those in the model explaining skewness. Finally, if we run a similar model with the median of the resting-state hitting-time distribution as the dependent variable, we only find a trend toward significance between schizophrenia and control in the opposite direction (*β* = 0.473, *t*(252) = 0.244, *p* = 0.054). Hierarchical sensory processing streams identified above as those with the longest hitting times show the largest changes between control and diagnosed groups, Fig. (9).

**Figure 9:**
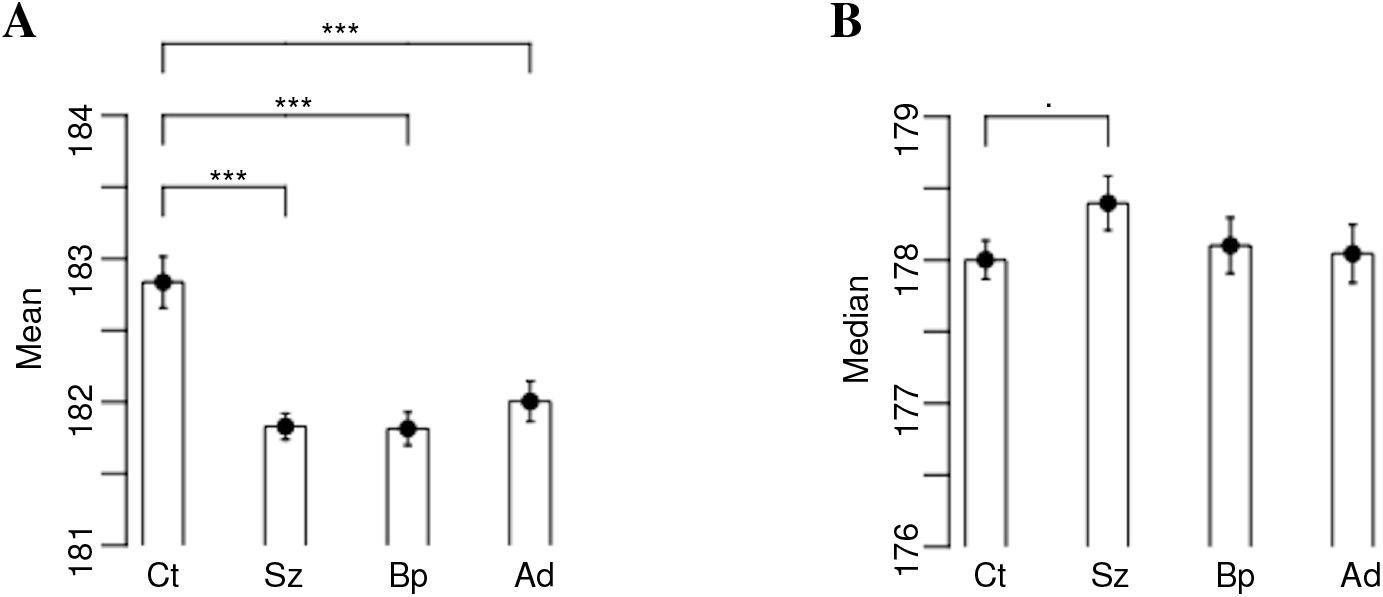
Mean of hitting-time distribution is significantly different across patient groups. Distribution (A) mean and (B) median of the hitting-time distribution in patient and control groups during resting-state scans. p-values significance codes:. = 0.1, * =< 0.05, ** =< 0.01, * * * =< 0.001.

## ACKNOWLEDGMENTS

Publication of this article was funded by the University of Colorado Boulder Libraries Open Access Fund. The research was supported by the Department of Electrical and Computer Engineering, University of California San Diego, and Department of Electrical, Computer and Energy Engineering and Institute of Cognitive Science of the University of Colorado Boulder. Piya Pal was supported in parts by NSF NCS-FO 1734940 and UC San Diego. We would also like to acknowledge members of Social Neuroscience and Games (SNaG) Lab, and Psychology and Neuroscience Department at the University of Colorado Boulder whose comments helped us improve our work. We would like to especially thank Heejung Jung, John Pearson, Terry Sejnowski, and David Smith for manuscript review during its preparation.

